# Efficient Cas9 multiplex editing using unspaced gRNA arrays engineering in a *Potato virus X* vector

**DOI:** 10.1101/2020.06.25.170977

**Authors:** Mireia Uranga, Verónica Aragonés, Sara Selma, Marta Vázquez-Vilar, Diego Orzáez, José-Antonio Daròs

## Abstract

Systems based on the clustered, regularly interspaced, short palindromic repeat (CRISPR) and CRISPR associated proteins (Cas) have revolutionized genome editing in many organisms, including plants. Most CRISPR-Cas strategies in plants rely on genetic transformation using *Agrobacterium tumefaciens* to supply the gene editing reagents, such as the Cas nucleases or the guide RNA (gRNA). While the Cas nucleases are constant elements in editing approaches, gRNAs are target-specific and a screening process is usually required to identify those most effective. Plant virus-derived vectors are an alternative for the fast and efficient delivery of gRNAs into adult plants, due to the virus capacity for genome amplification and systemic movement, a strategy known as virus-induced genome editing (VIGE). In this work, we engineered *Potato virus X* (PVX) to build a vector able to easily express one or more gRNAs in adult solanaceous plants. Using the PVX-based vector, *Nicotiana benthamiana* genes were efficiently targeted, producing nearly 80% indels in a transformed line that constitutively expressed *Streptococcus pyogenes* Cas9. Interestingly, results showed that the PVX vector allows expression of arrays of unspaced gRNAs achieving highly efficient multiplex editing in a few days in adult plant tissues. We also demonstrate that genome modifications are inherited in plants regenerated from infected tissues. In sum, the new PVX VIGE vector allows easy, fast and efficient expression of gRNAs arrays for multiplex CRISPR-Cas genome editing and will be a useful tool for functional gene analysis and precision breeding across diverse plant species, particularly in important crops of the family *Solanaceae*.

## Introduction

Targeted gene editing of plant DNA is a valuable tool for basic and applied biology as it facilitates gene function studies and crop improvement (Pennisi, 2010; Chen *et al.*, 2019). This can be performed by sequence-specific nucleases (SSNs) that specifically bind to the user-selected genomic region and induce DNA double-strand breaks (DSBs) (Voytas, 2013), but recent emergence of tools based on bacterial clustered, regularly interspaced, short palindromic repeat (CRISPR)–associated protein (Cas) systems have revolutionized targeted genome editing (Cong et al., 2013). Most common arrangements comprise the Cas9 endonuclease from *Streptococcus pyogenes* and a synthetic guide RNA (gRNA), which combines the functions of CRISPR RNA (crRNA) and trans-activating crRNA (tracrRNA). The gRNA directs the Cas9 endonuclease to a target sequence complementary to 20 nucleotides preceding the 5’-NGG-3’ protospacer-associated motif (PAM) required for Cas9 activity. The DSBs created by Cas9 activate the native host DNA repair mechanisms of non-homologous end-joining (NHEJ) or homology-directed repair (HDR). In plants, NHEJ occurs more frequently and results in small insertions or deletions (indels) that restore the integrity of the host DNA. These indels cause localized DNA disruption and have been used for sequence-specific knock-out of downstream gene products, such as proteins and long noncoding RNAs (Ran et al., 2013; Schiml et al., 2014). Thus, the specificity and versatility provided by the CRISPR-Cas tools allow for unprecedented, simple genome engineering of plant model species and economically important crops (Nekrasov *et al.*, 2013; Zhang *et al.*, 2016; Li *et al.*, 2017, 2018).

A key advantage of CRISPR genome editing consists in the possibility to program the cellular pool of Cas9 proteins with several gRNAs operating in parallel, a feature known as multiplexing. The multiplexing capacity of an editing tool determines the speed at which simultaneous modifications can be introduced in the genome and therefore the ability to perform comprehensive genome engineering. An efficient way to deliver several gRNAs acting simultaneously in the cell consists in engineering long RNAs comprising several gRNA units arrayed in tandem, which are later processed into single functional gRNAs. Contrary to other CRISPR nucleases as Cas12a (Zetsche *et al.*, 2017), Cas9 itself has no ability to process gRNA arrays on its own. Therefore, functional tandem gRNAs require processable spacers to be included in the array design. Spacers can be engineered involving either self-cleavable ribozymes (Gao and Zhao, 2014; Xu et al., 2017), or other RNA motifs processable by transacting RNases. For the second option, two main strategies have been described. One of them makes use of tRNAs as spacers, which are then removed by endogenous tRNA-processing RNases, RNase P and RNase Z (Xie et al., 2015). In the second approach, spacers are small recognition sequences processed by a trans-acting Cys4 RNase (Nissim et al, 2014; Tsai et al., 2014), which needs to be exogenously supplied, usually as a transgene.

CRISPR-Cas approaches in plants have mainly focused on the delivery of the nucleases and gRNAs by transformation technologies or transient delivery to protoplasts (Cong et al., 2013; Nekrasov et al., 2013). However, recent studies indicate that viral vectors may be most useful to express CRISPR-Cas reagents, as they are in other biological systems (Platt et al., 2014; Senís et al., 2014; Lau and Suh, 2017; Xu et al., 2019), following the so-called virus-induced genome editing (VIGE). Several plant RNA virus-based replicons have been tested as vectors for the delivery of gRNAs to create gene knockouts and insertions, including *Tobacco rattle virus* (TRV) (Ali *et al.*, 2015; Ellison *et al.*, 2020), *Tobacco mosaic virus* (Cody et al., 2017), *Pea early browning virus* (PEBV) (Ali et al., 2018), *Beet necrotic yellow vein virus* (BNYVV) (Jiang et al., 2019) and *Barley stripe mosaic virus* (BSMV) (Hu et al., 2019). Compared to delivery methods via *Agrobacterium tumefaciens*, plant virus-mediated gRNA delivery systems possess some advantages: (i) the gRNAs can accumulate to high levels owing to viral replication, which usually contributes to a higher genome editing efficiency; (ii) if the viral vector moves systemically, phenotypic alterations may appear in infected plants in a relatively short period after virus inoculation; and (iii) in the case of RNA viruses, they can produce a high incidence of edited cells without the risk of integration of heterologous material into the plant genome, which avoids raising additional regulatory and ethical issues. As drawbacks, each viral vector has its own particularities based on virus molecular biology and is restricted to a specific host range. Consequently, there is a necessity to enlarge and improve the available toolbox for VIGE.

*Potato virus X* (PVX) is a member of the genus *Potexvirus* (family *Alphaflexiviridae*) that infects 62 plant species of 27 families (Edwardson and Christie, 1997), including important crops in the family *Solanaceae*, such as potato, tomato, pepper or tobacco, and is transmitted mechanically from plant to plant (Adams et al., 2004). PVX has a plus (+) single-strand RNA genome 6.4-kb in length, with a 5’-methylguanosine cap, a polyadenylated 3’ tail, and five open reading frames (ORFs) (Loebenstein and Gaba, 2012). In this study, we describe a novel PVX-based gRNA delivery vector for CRISPR-Cas genome editing in plants, which produces targeted indels with high efficiency (nearly 80%) in *N. benthamiana* plants previously transformed to constitutively express Cas9 nuclease. We also demonstrate that PVX is a suitable vector for the simultaneous delivery of multiple gRNAs per viral genome. Unexpectedly, we found that, in the absence of spacers and processing signals, gRNAs engineered in tandem in the PVX genome are able to induce efficient gene editing. Efficiency of editing on unspaced gRNA arrays are apparently influenced by the relative position of each gRNA in the arrays, with positions close to 3’ end of the viral RNA precursor transcript showing higher efficiency. Finally, we demonstrate that whole plants carrying indels at the target genes can be efficiently regenerated from infected tissue, a strategy that could facilitate high throughput screenings.

## Results

### Engineering a PVX-based gRNA delivery system for gene editing in plants

CRISPR-Cas9 is a two-component system that requires both Cas9 nuclease and a small gRNA to perform gene editing. We aimed to develop a gRNA delivery system based on PVX, an RNA virus that infects many plant species, particularly important crops in the family *Solanaceae*. To set up this system, we used an *N. benthamiana* transformed line that constitutively expresses a version of *Streptococcus pyogenes Cas9* (SpCas9) under the control of *Cauliflower mosaic virus* (CaMV) 35S promoter and *A. tumefaciens Nopaline synthase* (nos) terminator (Bernabé-Orts et al., 2019). More specifically, this line overexpresses a human codon-optimized version of SpCas9 coding sequence fused to a nuclear localization signals (NLS) at the carboxyl terminus, which have been shown to improve Cas9 activity in plants (Yin et al., 2015). Our PVX-based gRNA delivery system (PVX::gRNA) was designed using as a template a PVX mutant where the 29 initial codons of the coat protein (CP) are deleted and heterologous RNA expression is controlled by the subgenomic CP promoter. In turn, PVX CP is expressed from an heterologous promoter derived from that of CP of *Bamboo mosaic virus* (BaMV; genus *Potexvirus*). The PVX::gRNA system was constructed by cloning a cDNA corresponding to the 96-nt gRNA downstream the PVX CP promoter. The gRNA consisted of a 20-nt protospacer sequence specific to the target gene and a 76-nt scaffold highly conserved in the CRISPR/Cas9 system, also known as direct repeat.

Xie et al. (2015) showed that the host endogenous tRNA-processing system cleaves tandemly arrayed tRNA-gRNA constructs into gRNAs, thus improving the efficiency of gene editing. Aiming to explore whether this strategy also applies to splice out the gRNA from the viral subgenomic RNA that is transcribed from the BaMV CP promoter, four different constructs were designed: (i) gRNA flanked with tRNAs at both 5’ and 3’ ends (tR-gRNA-tR); (ii) gRNA flanked with tRNA only at 5’ end (tR-gRNA); (iii) gRNA flanked with tRNA only at 3’ end (gRNA-tR); and (iv) gRNA without any flanking tRNAs (gRNA) (Figure 1).

**Figure 1.**
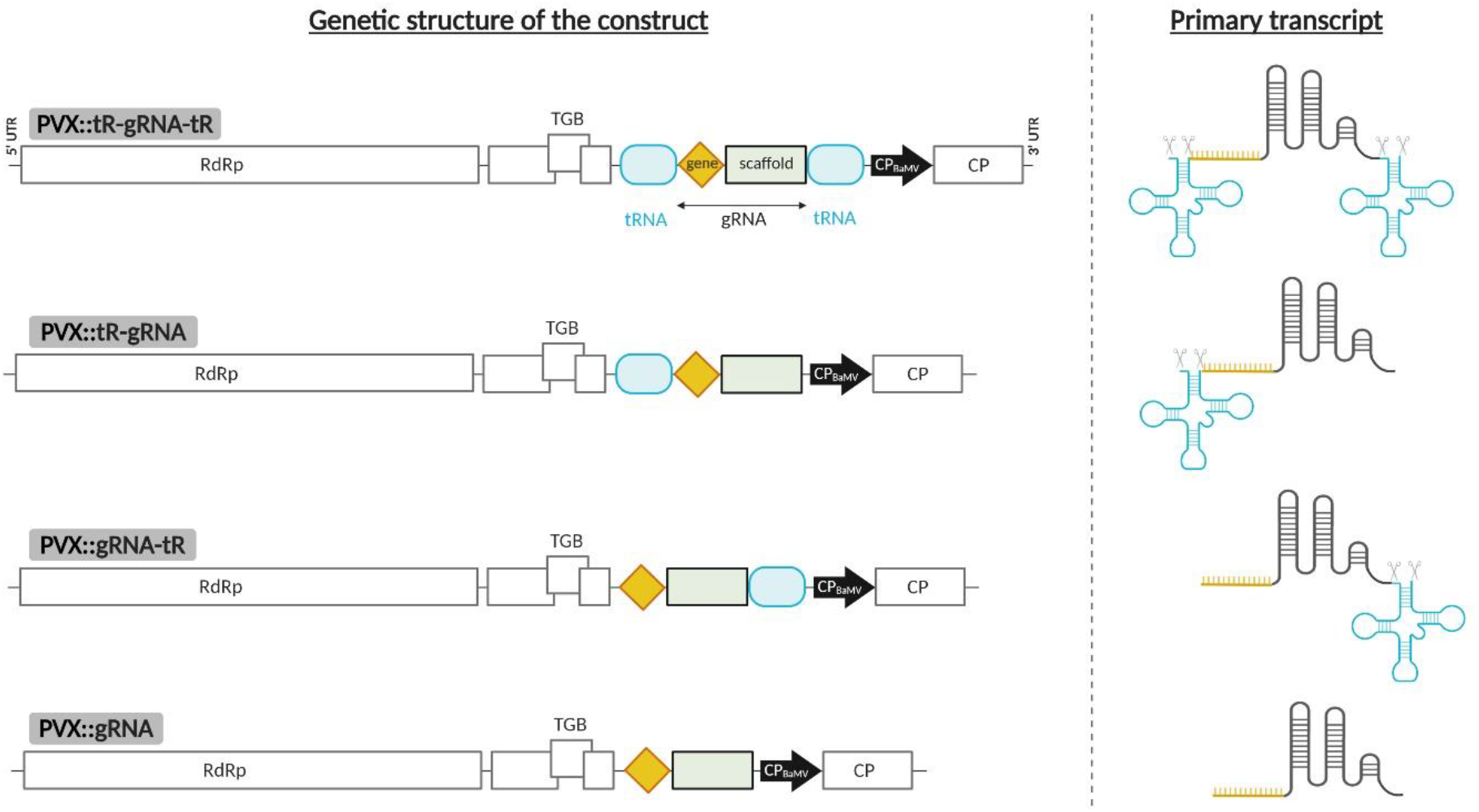
PVX vector to express gRNAs for CRISPR-Cas9-based gene editing in *N. benthamiana*. Left panel, schematic representation of recombinant clones PVX::tR-gRNA-tR, PVX::tR-gRNA, PVX::gRNA-tR and PVX::gRNA, which express different versions of a gRNA flanked or not with tRNAs to promote RNA processing by endogenous RNases. Right panel, structure of the subgenomic primary transcripts of each PVX recombinant clone. RdRp, RNA-dependent RNA polymerase; TGB, triple gene block; and CP, coat protein, are represented by open boxes. Heterologous *Bamboo mosaic virus* CP promoter (CP_BaMV_) is represented by a black arrow. 5′ and 3′ untranslated regions (UTRs) are represented by lines. tRNA sequences are represented by round, light blue boxes. gRNA consists of a gene-specific 20-nt protospacer (orange diamond) and a conserved 76-nt scaffold (grey box). CP_BaMV_, tRNAs, protospacer and scaffold, not at scale.

### PVX::gRNA system can efficiently target host genes without tRNA-mediated processing of the gRNA

As a proof-of-concept of the PVX::gRNA system, the *NbXT2B* locus was selected as the target gene to be edited. Previous work assessed the efficiency of Cas9-mediated gene editing of several *N. benthamiana* loci, including this gene, with different gRNAs (Bernabé-Orts et al., 2019). XT2B is a xylosyl transferase, and loss of function of this gene alters the glycosylation pattern in endogenous and recombinant proteins (Cavalier and Keegstra, 2006). Thus, *NbXT2B*-specific gRNA was cloned into the four different tRNA-gRNA constructs mentioned above to generate the PVX::tR-gXT2B-tR, PVX::tR-gXT2B, PVX::gXT2B-tR and PVX::gXT2B derivatives (Figure 2a). *A. tumefaciens* carrying the corresponding plasmids were then inoculated into leaves of Cas9 *N. benthamiana*. As a control for gene editing, additional plants were inoculated with *A. tumefaciens* carrying PVX::crtB. This is a visual tracker of PVX infection and movement previously developed, which results in a bright yellow pigmentation of infected tissue (Majer et al., 2017).

**Figure 2.**
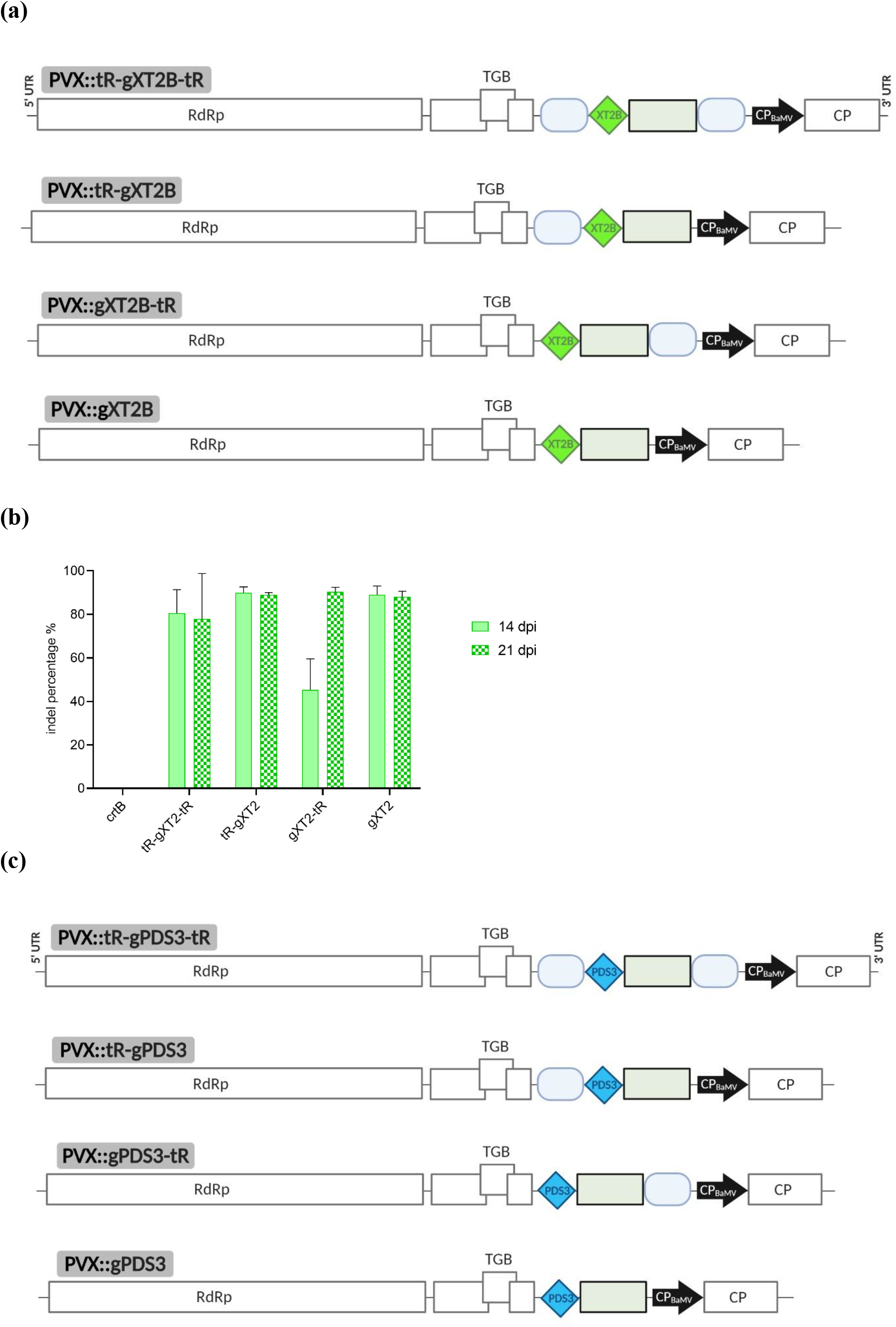
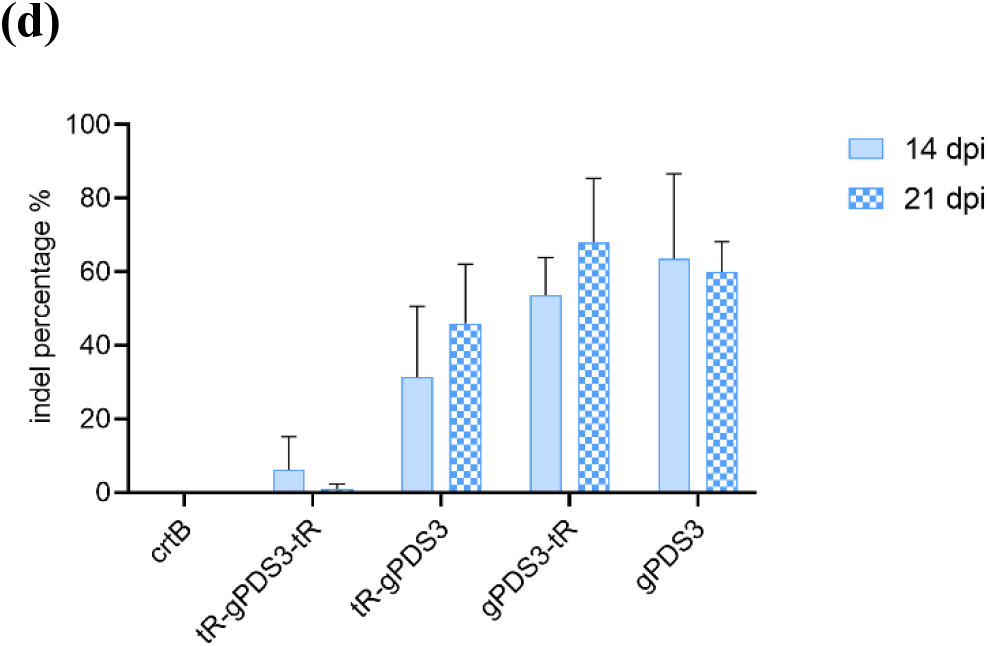
Single gene editing, with or without flanking tRNAs, using the PVX-based vector in *N. benthamiana*. (a) Schematic representation of recombinant clones PVX::tR-gXT2B-tR, PVX::tR-gXT2B, PVX::gXT2B-tR and PVX::gXT2B. *NbXT2B* protospacer is represented by a green diamond. Other details as in the legend to Figure 1. (b) Inference of CRISPR edits (ICE) analysis of the first systemically infected upper leaf of *N. benthamiana* plants inoculated with PVX::gXT2B derivatives at 14 (n=3) and 21 dpi (n=3). PVX::crtB was used as a negative editing control. (c) Schematic representation of recombinant clones PVX::tR-gPDS3-tR, PVX::tR-gPDS3, PVX:: gPDS3-tR, PVX::gPDS3 and PVX::gPDS3. *NbPDS3* protospacer is represented by a blue diamond. Other details as in the legend to Figure 1. (d) ICE analysis of the first systemically infected upper leaf of *N. benthamiana* plants inoculated with PVX::PDS3 derivatives at 14 (n=3) and 21 dpi (n=3). (b and d) Columns and error bars represent average indels (%) and standard deviation, respectively.

At 7 days post-inoculation (dpi), symptoms of PVX infection characterized by the appearance of vein banding, ring spots and leaf atrophy were observed in the upper non-inoculated leaves of all PVX::gRNA infiltrated *N. benthamiana* plants. These symptoms persisted and became more noticeable over time. However, symptoms of PVX::gXT2B were slightly more intense than those of PVX::tR-gXT2B, PVX::gXT2B-tR, and at the same time, symptoms of these two last virus were slightly more intense than those of PVX::tR-gXT2B-tR. Samples from the first systemically infected upper leaf were collected at 14 and 21 dpi. On the one hand, RNA was extracted and reverse transcription (RT)-polymerase chain reaction (PCR) analysis confirmed the presence of PVX in all inoculated plants. The insertion of the tRNA-gRNA region in the PVX progeny was also confirmed by RT-PCR. These results indicated that, although addition of one or two tRNA sequences into the genome may be affecting virus fitness, they do not abolish PVX infectivity and systemic spread, and that the heterologous tRNA-gRNA region is conserved in PVX progeny over time. On the other hand, DNA was extracted from the leaf samples collected at 14 and 21 dpi and a 750-bp fragment of the *NbXT2B* gene covering the Cas9 target site was amplified by PCR. Sanger sequencing of the PCR products and inference of CRISPR edits (ICE) analysis revealed an efficient genome editing in systemic leaves of all PVX::gRNA inoculated plants (Figure 2b). No statistically significant differences were observed in *NbXT2B* gene editing among the four different PVX::gRNA derivatives, with average indel percentage ranging from 37% to 85%. These results suggest that all PVX::gRNA derivatives are suitable for targeted editing of *N. benthamiana* genes in the presence of Cas9, regardless of the addition of a tRNA sequence at 5’ and/or 3’ ends of the gRNA. To confirm this finding, the experiment with the four PVX::gRNA derivatives was repeated following the same procedure, but this time selecting *NbPDS3* as the target gene (Figure 2c). Phytoene desaturase (PDS) is a key enzyme of the carotenoid biosynthetic pathway in plants (Burch-Smith et al., 2006). At 14 and 21 dpi, the first systemically infected upper leaf was sampled, following extraction of genomic DNA and PCR amplification of a 650-bp fragment covering the Cas9 target site of *NbPDS3*. For this gene, ICE analysis displayed no editing for PVX::tR-gPDS3-tR, a relatively low efficiency (25%) for PVX::tR-gPDS3, and high efficiency (68-73%) for PVX::gPDS3-tR and PVX::gPDS3 (Figure 2d). Owing to its high efficiency and design simplicity, PVX::gRNA constructs lacking tRNA sequences were used for subsequent experiments.

### Efficient and easy editing of multiple genes from a single PVX construct

After the successful gene editing of endogenous genes in *N. benthamiana*, we wondered whether our PVX-based vector could be feasible for the delivery of multiple gRNA molecules from a single construct (i.e. multiplexing). The capacity to edit several genes at once using transient gRNA delivery strategies remains of great interest for generating plants with multiple gene knockouts. In addition to the *N. benthamiana* genes tested through the single gRNA strategy (*NbXT2B* and *NbPDS3*), we decided to include an additional target gene for the multiplexing. Based on the results on editing efficiency obtained by Bernabé-Orts et al. (2019), we selected *NbFT*. Flowering locus T (FT) promotes the development of inflorescences from vegetative meristem, and loss of function of this gene results in late flowering (Wigge et al., 2005). Considering the finding that our PVX::gRNA system can efficiently produce indels on target genes without any gRNA processing, we hypothesized that multiple gRNAs could be tandemly delivered within a single PVX construct. We further proposed to position the three gRNA sequences next to each other, without any spacer sequence and under the control of the same BaMV CP promoter. Thus, using the PVX vector as a template, gPDS3 and gXT2B were positioned on the 5’ and 3’ ends of the gRNA construct, respectively, and gFT was included between them, creating the PVX::gPDS3:gFT:gXT2B construct (Figure 3a). Additionally, a PVX::gFT construct harbouring the single gRNA for *NbFT* was designed to compare the editing efficiency between single and multiplex gRNA strategies (Figure 3a). *A. tumefaciens* carrying either PVX::gPDS3:gFT:gXT2B or PVX::gFT were inoculated into Cas9 *N. benthamiana* leaves, and inoculation with PVX::crtB was used as a control for gene editing. Again, at 7 dpi symptoms of PVX infection were observed in the upper non-inoculated leaves of all PVX::gRNA infiltrated *N. benthamiana* plants. Samples from the first systemically infected upper leaf were collected at 7, 14, 21 and 28 dpi.

**Figure 3.**
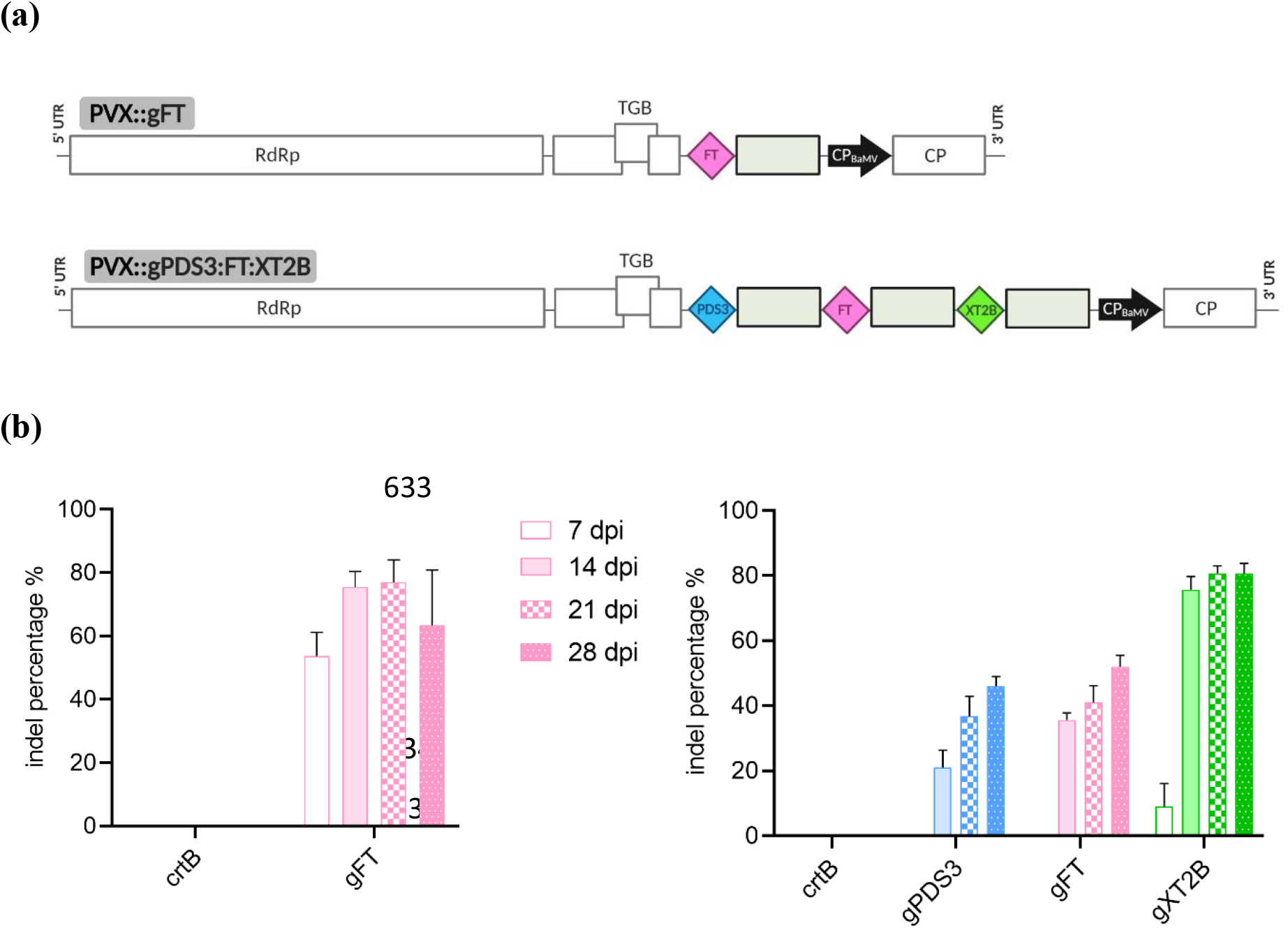
Single versus multiplex gene editing using the PVX-based system in *N. benthamiana*. (a) Schematic representation of recombinant clones PVX::gFT (single gRNA strategy) and PVX::gPDS3:gFT:gXT2B (multiplex gRNA strategy). Protospacers for *NbPDS3, NbFT* and *NbXT2B* are represented by blue, pink and green diamonds, respectively. Other details as in the legend to Figure 1. (b) ICE analysis of the first systemically infected upper leaf of *N. benthamiana* plants (n=4) inoculated with PVX::gFT (left) and PVX::gPDS3:gFT:gXT2B (right). PVX::crtB was used as a negative control. dpi, days post-inoculation. Columns and error bars represent average indels (%) and standard deviation, respectively.

Following DNA extraction and PCR amplification of the target sites, *NbPDS3*, *NbFT* and *NbXT2B* amplicons were subjected to ICE analysis (Figure 3b). For PVX::gPDS3:gFT:XT2B infected leaves, at 7 dpi indels were detected only in *NbXT2B* gene but at a low percentage (9%). Remarkably, at 14 dpi gene editing was boosted as indels were detected in all three genes. However, efficiencies were determined by gRNA positioning on the heterologous construct: the lowest average indel percentage (21%) corresponded to gPDS3 that was located in the 5’ side of the gRNA precursor; gFT, located in the middle of the precursor, showed 36% indels; whereas the highest indel percentage (75%) corresponded to gXT2B, located in the 3’ side. A slight increase in gene editing was observed over time for *NbPDS3* and *NbFT* (Figure 3b, right panel). Time-course comparison with the single gRNA construct, in the case of gFT, showed that multiplexing lowers the editing efficiency (Figure 3b, compare left and right graphs). These results indicate that various gRNAs can be delivered simultaneously using the PVX-based vector, although indel production is lower compared to single gRNA delivery. Moreover, results suggest that editing efficiency is dependent on gRNA positioning on the heterologous construct, being the gRNA proximal to the 3’ end the most efficient among all.

### Editing efficiency is increased in plants regenerated from tissue infected with PVX::gRNA vector

The above-explained results demonstrate that the PVX-based vector is suitable for the delivery of either single or multiple gRNAs leading to efficient editing of target genes. It is well known that PVX is transmitted mainly by mechanical contact between infected and healthy plants, although transmission by zoospores of the fungus *Synchytrium endobioticum* has also been reported (Loebenstein and Gaba, 2012). Moreover, it is well established that that this virus is not transmitted through true seed or pollen. As *NbXT2B* appeared to be the most efficiently edited gene, seeds from PVX::gXT2B infected plants were recovered. The progeny was then screened for the presence of PVX and heritability of editing in *NbXT2B*. All plants were virus-free, but none of them carried indels at the target site of *NbXT2B*. These results indicate that PVX is unable to infect germline cells and therefore the progeny of transiently modified plants cannot inherit sequence modification in the target genes.

We proposed that leaf tissue from plants infected with PVX::gRNA could be regenerated into whole plants and screened for the presence of gene editing. We decided to test this possibility for plants modified at either single or multiple target genes. Thus, leaf discs from plants inoculated with PVX::gXT2B or PVX::gPDS3:gFT:gXT2B were collected at 14 and 21 dpi and regenerated following tissue culture. Next, we studied whether the regenerated plants, which still showed symptoms of virus infection, carried modifications in the target genes. ICE analysis of PCR products indicated a strong presence of indels in plants regenerated from leaf tissue that had been edited at either single or multiple genes, regardless of the sampling time (Figure 4). Surprisingly, editing efficiency in regenerated plants was higher compared to that of parental tissues: from 77% to 95.8% for *NbXT2B* in single gRNA delivery strategy (Figure 4, left panel); and from 46% to 74% for *NbPDS3*, from 52% to 78.5% for *NbFT*, and from 52% to 94.5% for *NbXT2B* in multiple gRNA delivery strategy, respectively (Figure 4, right panel).

**Figure 4.**
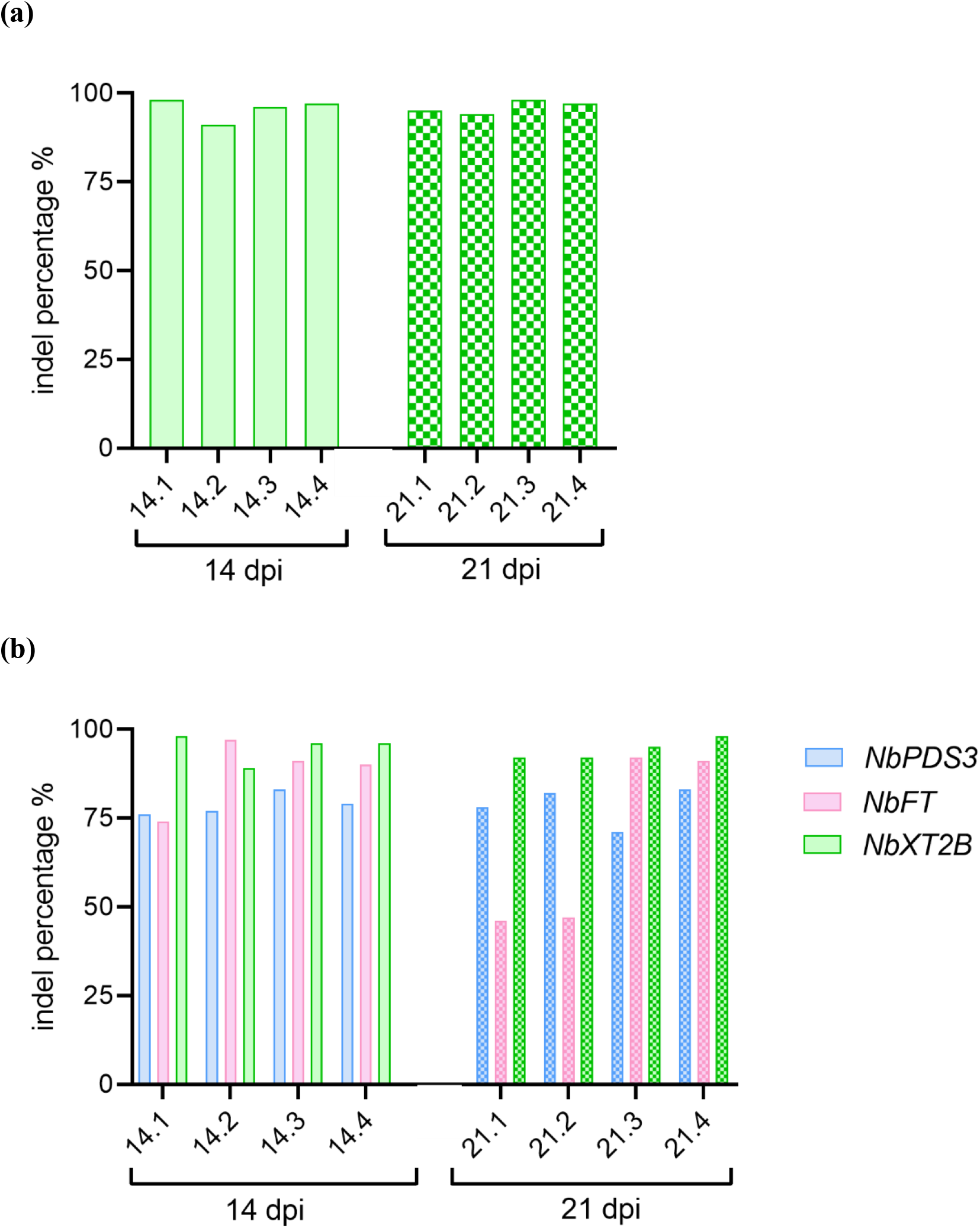
Gene editing in *N. benthamiana* plants regenerated from PVX-infected tissue. ICE analysis of plants regenerated from leaf discs infected with (a) PVX::gXT2B and (b) PVX::PDS3:FT:XT2B. Four independent plants regenerated from leaf discs collected at 14 dpi or 21 dpi, as indicated, are represented as 14.1-14.4 or 21.1-21.4, respectively. Columns represent average indels (%).

## Discussion

Targeted editing of plant genomes by CRISPR-Cas9 technology is expected to play a crucial role in both basic genetic studies and crop improvement for 21^st^-century agriculture. In this context, the use of plant viral vectors for the transient delivery of CRISPR-Cas9 components has emerged as a promising strategy over conventional transformation technologies. The first VIGE reports focused on generation of gene knockout plant lines using gRNA delivery systems based on geminiviruses (Baltes et al., 2014; Yin et al., 2015). Subsequent studies led to the development of several plant RNA virus-based vectors with the same purpose (Ali *et al.*, 2015, 2018; Cody *et al.*, 2017; Hu *et al.*, 2019; Jiang *et al.*, 2019; Ellison *et al.*, 2020). Here, we describe a PVX-based gRNA delivery vector that allows easy multiplexing for efficient targeted editing of the model species *N. benthamiana* as a novel approach to expand the current VIGE toolbox.

First, we explored the potential of PVX for the transient delivery of gRNA molecules into host plants. Several studies have documented that nucleotide overhangs on either 5’ or 3’ ends of the gRNA negate Cas9 activity *in vitro*, which leads to a reduction of DSBs (Jinek et al., 2012; Mali et al., 2013; Dahlman et al., 2015; Zalatan et al., 2015). Considering that the host endogenous tRNA-processing system can splice out tandemly arrayed tRNA-gRNA constructs into mature gRNAs (Xie et al., 2015), we designed different PVX::gRNA derivatives where the gRNA was flanked or not with tRNAs (Figure 1). We hypothesized that, although host plant tRNA processing machinery cleaves some genomes, enough virus will survive to maintain infection. Of note, pre-tRNA processing occurs in the nucleus and PVX replication in the cytoplasm. Certainly, the presence of one or two tRNA sequences in the viral genome had a minor effect on PVX infectivity and the ability to move systemically. Surprisingly, we also found that long gRNA overhangs at both ends did not affect Cas9 catalytic activity *in planta*, as there were no substantial differences on editing efficiency of target genes among the PVX::gRNA derivatives with no tRNA and those with tRNAs at either 5’ or 3’ side (Figures 2b and 2d). The capacity to produce biologically active gRNAs without tRNA-mediated processing follows findings from previous VIGE reports in which gRNAs contained long 3’ extensions (Cody et al., 2017; Ali et al., 2018). Even more surprising is the observed ability of unspaced tandem gRNA arrays to drive efficient editing, a feature earlier observed by Cody et al. (2017) in TMV-based VIGE experiments using a smaller tandem comprising only two gRNAs, and very recently also showed in a TRV-based system (Ellison *et al.*, 2020). To our knowledge, no other biological system has been described where Cas9 shows this level of permissiveness to the presence of 5’ prime extensions, neither are we aware of other systems where spacer-less gRNA arrays are fully functional for gene editing. These findings suggest that one or more of the following processes may be occurring *in vivo*: (i) Cas9 tolerates large, non-complementary overhangs within the gRNA protospacer (5’ end) *in planta*; (ii) Cas9 is able to precisely cleave gRNA overhangs *in planta*; (iii) endogenous RNases cleave gRNA overhangs *in planta;* or (iv) the virus, as result of imperfect replication or partial RNA degradation, produces a relatively large population of subgenomic fragments some of which, by chance, contain the correct 5’ ends. Since the functionality of unspaced gRNAs in Cas9 multiplexing has been only reported associated to viral vectors, we favour the last explanation, perhaps in combination with other mechanisms. For instance, direct repeats may protect gRNA scaffold from degradation by unspecific plant RNases, favouring unspecific cleavage to take place in and around the protospacer and therefore increasing the relative concentration of correctly cleaved (functional) gRNA species. Regardless of the precise mechanism underlying native gRNA processing, our results reinforce the fact that gRNAs can be launched from viral vectors without the need for processing spacer elements, which rather simplifies construct design.

One of the main advantages of VIGE is that high gRNA levels are accumulated into host plants due to virus replication which leads to a fast and efficient editing process not achievable with conventional delivery methods (Cody and Scholthof, 2019). In fact, we observed a high editing efficiency when single gRNAs were delivered through PVX, reaching nearly 80% of indels (Figure 2b, and Figure 3b, left panel). Remarkably, when multiple gRNAs were delivered within the same PVX vector with no processing spacers between gRNAs, some bias was observed related to gRNA positioning on the construct. gRNA proximity to the 5’ end of the viral RNA apparently caused a decrease in indel production, while when gRNA was located on the 3’ end, the indel rates were comparable to those observed for single gRNA delivery. This finding may be related with the mechanism of gRNA processing in viral vectors discussed above and will require further analysis.

The capacity to transmit genome modifications to the next generation is considered a desired feature of any VIGE system. Starting from infected tissue where single or multiple gRNA had been delivered, we regenerated whole plants and screened them for the presence of modifications. The progeny showed an increase on indel percentages compared to those observed on parental tissue, and most importantly, all of them carried biallelic mutations for *NbXT2B* gene (Figure 4). This confirmed that genome editing is inherited to subsequent generations. However, regenerated plants developed symptoms of viral infection, indicating that PVX was still present. It is well documented that PVX is unable to infect germline cells and therefore is not transmitted through true seed or pollen (Loebenstein and Gaba, 2012). Therefore, virus-free edited plants can be obtained in the T1 generation from seeds of regenerated plants. This is also the stage where Cas9 is usually segregated in traditional transgenesis approaches. Therefore, PVX-based VIGE offers similar speed than transgenics to obtain transgene-free edited plants, but a higher efficiency and now also multiplexing capacity. Combining heat treatment before the excision of edited leaves and the incorporation of ribavirin into the culture medium have also been reported to enhance the proportion of virus-free progeny (Faccioli, 2001). Importantly, a recently published VIGE strategy, based on fusion of RNA sequences that promote cell-to-cell mobility, has shown to accelerate the recovery of mutant progeny (Ellison *et al.*, 2020). In this respect, it should be noticed that PVX-gRNAs can be a very useful tool when combined with transgenics to enhance multigene editing efficiency. Often, multiplex editing with a transgenic construct miss some of the intended targets due to our limited capacity to predict gRNAs efficiency. Provided that Cas9 is already present in T0 lines, if missing targets are early genotyped in these lines, they could then be infected with multiplex PVX-gRNAs designed specifically to fill the editing gaps. This approach can be faster and more efficient than re-transformation, and most importantly, it circumvents the need for a second selection marker.

In summary, the virus-mediated genome editing system described here postulates PVX as a promising vector for the delivery of one or more gRNAs without the need of processing elements, thus leading to a highly efficient gene editing without the risk of integration of viral sequences into the plant genome. These modifications on the target genes are inherited to the next generation when plants are regenerated from infected tissue. Furthermore, the wide range of host species that PVX can infect, including important crops in the family *Solanaceae*, highlights its potential for CRISPR-Cas9-based modification of economically relevant species. In addition, PVX is a preferred vector in plant biotechnology (Röder et al., 2019). Our findings expand the current knowledge about the VIGE toolbox and will contribute to future applications in plant functional genomics and agricultural biotechnology.

## Experimental procedures

### Design of gRNAs and vector construction

Three *N. benthamiana* genes were chosen as targets for CRISPR-Cas9 mediated gene editing: *Flowering locus T* (*NbFT*), *Phytoene desaturase 3* (*NbPDS3)* and *UDP-xylosyltransferase 2* (*NbXT2B*). The design of gRNAs was performed using the CRISPR-P online tool as described in Bernabé-Orts et al. (2019). Target sequences for each gene are described in Table S1. Plasmid pPVX contains a PVX UK3 strain infectious cDNA flanked by the *Cauliflower mosaic virus* (CaMV) 35S promoter and *A. tumefaciens Nopaline synthase* (nos) terminator. In all PVX-based recombinant clones, heterologous genes were expressed from PVX coat protein (CP) promoter, and PVX CP was expressed from an heterologous promoter derived from *Bamboo mosaic virus* (BaMV; genus *Potexvirus*). T 29 initial codons of PVX CP were deleted (Dickmeis et al., 2014). Plasmids to express recombinant viruses PVX::tR-gXT2B-tR, PVX::tR-gXT2B, PVX::gXT2B-tR, PVX::gXT2B, PVX::tR-gPDS3-tR, PVX::tR-gPDS3, PVX::gPDS3-tR, PVX::gPDS3, PVX::gFT; and PVX::gPDS3:gFT:gXT2B were built by standard molecular biology techniques, including PCR amplification of target gRNAs with high-fidelity Phusion DNA polymerase (Thermo Scientific) and Gibson DNA assembly with the NEBuilder HiFi DNA assembly master mix (New England Biolabs). Primers used for vector construction are listed in Tables S2 and S3. Recombinant virus PVX::crtB that expresses *Pantoea ananatis* phytoene synthase (crtB), which induces a distinctive yellow pigmentation in infected tissue (Majer et al., 2017), was used as a control for the experiments. The sequences of all the recombinant PVX-derived clones were confirmed by standard DNA sequencing techniques and are included in Figure S1.

### Plant growth conditions and inoculation

Transgenic Cas9-overexpressing *N. benthamiana* plants (Figure S2) were grown in growth chambers at 25°C under a 16-h-day/8-h-night cycle. Fully expanded upper leaves from 4- to 6-week-old plants were used for inoculation of PVX::gRNA constructs. Electrocompetent *A. tumefaciens* C58C1 containing the helper plasmid pCLEAN-S48 (Thole et al., 2007) were transformed with plasmids containing the different PVX recombinant clones. Transformed cells were spread on to 50 μg/L kanamycin, 50 μg/L rifampicin and 12.5 μg/L tetracycline Luria Bertani (LB) agar plates. Single colonies were grown overnight at 28°C in 10 mL LB containing 50 μg/mL kanamycin. At optic density at 600 nm (OD_600_) 1-2, cells were pelleted by centrifuging at 7800 revolutions per min (rpm) for 5 min and resuspended to an OD_600_ of 0.5 in infiltration buffer (10 mM 2-(N-morpholino)ethanesulfonic acid (MES)-NaOH, 10 mM MgCl_2_ and 150 μM acetosyringone, pH 5.6) (Bedoya et al., 2010). Resuspended bacteria were incubated at 28°C for 2 h. Two leaves per plant were agroinfiltrated on the abaxial side using a 1-mL syringe. Control plants were inoculated with PVX::crtB following the same procedure. Immediately following agroinfiltration, plants were watered and transferred to a growth chamber under a 12-h-day/12-h-night and 25°C/23°C cycle. Samples from the first symptomatic systemic leaf were collected with a 1.2-cm cork borer (approximately 100 mg of tissue), immediately frozen in liquid nitrogen and stored at −80°C until use.

### RT-PCR analysis of PVX progeny

Leaf samples were ground with a VWR Star-Beater for 1 min at 30 s^−1^, homogenized in extraction buffer (4 M guanidinium thiocyanate, 0.1 M sodium acetate, 10 mM EDTA and 0.1 M 2-mercaptoethanol, pH 5.5), and centrifuged at 13,000 rpm for 5 min. Next, the collected supernatants were mixed with 0.65 volumes of 96% ethanol and centrifuged at 13,000 rpm for 1 min. The remaining steps for RNA purification were performed using silica gel columns (Zymo Research) that were finally eluted with 10 μL of 10 mM Tris-HCl pH 8.5. 1 μL RNA was subjected to reverse transcription with RevertAid reverse transcriptase (Thermo Scientific) using primer D2409 in a 10-μL volume reaction, and 1 μL of this reaction was subjected to PCR amplification with *Thermus thermophilus* (*Tth*) DNA polymerase (Biotools B&M) using gene-specific primers in a 20-μL volume reaction. Reaction conditions were as previously described (Bedoya et al., 2010). Primers D2410 and D3436 were designed to amplify a 627-nt region of PVX genome corresponding to CP ORF. Primers D1789 and D2069 were designed to amplify the heterologous gRNA region. Primers used for RT and RT-PCR are listed in Table S4. RT-PCR products were analyzed by electrophoresis in 1% agarose gels in TAE buffer (40 mM Tris, 20 mM sodium acetate, and 1 mM EDTA, pH 7.2) and staining with ethidium bromide.

### Detection of Cas9-mediated gene editing

To purify DNA from leaf samples, the same procedure explained above for RNA was followed, except for not adding ethanol to the extract before loading the silica gel column. To confirm CRISPR/Cas9-mediated editing of the target genes, products covering *NbFT, NbPDS3* and *NbXT2B* target sites were obtained by PCR amplification of 1 μL genomic DNA using high-fidelity Phusion DNA polymerase and gene-specific primers. The resulting PCR products were purified using silica gel columns after separation by electrophoresis in 1% agarose gels and subjected to Sanger sequencing. The presence of sequence modifications was analyzed using the inference of CRISPR edits (ICE) software (https://www.synthego.com/products/bioinformatics/crispr-analysis). Primers used for amplification of target genes and sequencing are listed in Table S5.

### *In vitro* regeneration of edited *N. benthamiana* plants

Leaves from 4- to 5-week-old *N. benthamiana* plants were infiltrated with *A. tumefaciens* C58C1 cultures containing plasmids with the PVX recombinant clones PVX::gXT2B and PVX::gPDS3:gFT:gXT2B. As a control for gene editing, additional plants were infiltrated with PVX::crtB. At 14 and 21 days post-inoculation (dpi), the first upper leaf showing symptoms of viral infection was removed from each plant and surface-sterilized by submersion in sterilization solution (10% bleach, 0.02% Nonidet P-40) for 10 min. Leaves were then washed three times in sterile water. Sterilized leaves were cut into discs using a 1.2-cm cork borer and plated onto regeneration media (1X Murashige and Skoog (MS) with vitamins Gamborg B5, 0.5 g/L MES, 20 g/L sucrose, 8 g/L phytoagar, 1 mg/L 6-benzylaminopurine (6-BAP), 0.1 mg/L 1-naphthaleneacetic acid (NAA) and 50 μg/mL kanamycin, pH 5.7). Leaf discs were transferred to fresh plates every 2 weeks until shoots emerged. Shoots were cut and transferred to root induction media (1X MS with vitamins, 0.5 g/L MES, 10 g/L sucrose, 6 g/L phytoagar, 0.1 mg/L NAA and 50 μg/mL kanamycin, pH 5.7). Regenerated plantlets with roots were transferred into soil and covered with plastic cups to keep moisture. After 2 weeks, plastic cups were removed for plant normal growth.

## Accession numbers

Niben101Scf01519g10008.1 (FT); Niben101Scf01283g02002.1 (NbPDS3) Niben101Scf04551g02001.1 (XT2B);

## Acnowledgements

This work was supported by grants BIO2017-83184-R and BIO201678601-R from Ministerio de Ciencia e Innovación (Spain; co-financed European Union ERDF). M.U. is the recipient of the FPU17/05503 fellowship from Ministerio de Ciencia e Innovación (Spain).

## Supporting information

**Figure S1** Full sequence of wild-type Potato virus X (PVX) and its derived recombinant viruses PVX::crtB, PVX::tR-gXT2B-tR, PVX::tR-gXT2B, PVX::gXT2B-tR, PVX::XT2B, PVX::tR-gPDS3-tR, PVX::tR-gPDS3, PVX::gPDS3-tR, PVX::PDS3, PVX::gFT and PVX::gPDS3:gFT:gXT2B.

**Figure S2** Sequence of the transgenes inserted into *Nicotiana benthamina* to generate a line that constitutively expresses a human codon-optimizdd *Streptococcus pyogenes* Cas9.

**Table S1** *N. benthamiana* genes targeted by the PVX-based CRISPR/Cas9 system. PAMs are highlighted on grey background.

**Table S2** Primers used for the construction of recombinant viruses.

**Table S3** Primer combinations used for the construction of recombinant viruses.

**Table S4** Primers used for PVX diagnosis by RT-PCR.

**Table S5** Primers used for Cas9-gRNA gene editing analysis.

